# Forecasting the adoption and spread of a community-based marine management initiative using agent-based models

**DOI:** 10.1101/2024.06.16.599026

**Authors:** Andreas Christ Sølvsten Jørgensen, Thomas Pienkowski, Matthew Clark, Mathilda Dunn, Arundhati Jagadish, Alvaro Roel Bellot, Sangeeta Mangubhai, Alifereti Tawake, Margaret Tabunakawai Vakalalabure, Elisabeta Waqa, Tanya O’Garra, Hugh Govan, Vahid Shahrezaei, Morena Mills

## Abstract

While many successful initiatives for conserving nature exist, efforts to take them to scale have been inadequate. Moreover, conservation science currently lacks a systematic methodology for determining if or when interventions will reach effective scales and how programmatic decisions will affect the scaling process. This paper presents a modelling framework that aims to address both issues by operationalizing Diffusion of Innovations theory and local knowledge using agent-based modelling and Bayesian inference. By applying our framework to existing data on the spatiotemporal adoption of a community-based marine management initiative in Fiji, we demonstrate that our approach can identify the mechanisms that govern the observed adoption patterns. In this case, the relative advantage of the intervention, village social networks, and perceived knowledge stand out as important drivers of adoption. Using the identified causal processes, our approach can forecast business-as-usual and counterfactual future scenarios and hence inform conservation policy. Finally, we highlight the importance of spatiotemporal data for making detailed scaling predictions. We structure the paper as a step-by-step guide, highlighting our modelling decisions and possible limitations. Thus, besides presenting a case study, this paper serves as a template for practitioners and researchers to better model the scaling process of other conservation interventions.

## Introduction

Humanity faces the triple challenge of reversing biodiversity loss and limiting climate change while enhancing human well-being [1]. These challenges are substantial, with 25 per cent of assessed plant and animal species threatened with extinction [2], global temperatures projected to rise above 2°C [3], and seven of the eight ‘safe and just Earth system boundaries’ currently exceeded [4]. To address these growing crises, the United Nations has instituted three interconnected global policy frameworks: the Convention on Biological Diversity [5], the Framework Convention on Climate Change, and the Agenda for Sustainable Development. But despite bright spots of local success [e.g., 6, 7], global progress toward these coupled goals has been inadequate [8]. For instance, none of the CBD’s 20 Aichi Biodiversity Targets were met by their target date of 2020 [5]. Yet, there have been widespread commitments to scale conservation actions to ‘bend the curve’ upwards on biodiversity loss and curb climate change [9]. For example, Target 3 of the CBD Kunming-Montreal Global Biodiversity Framework aims to nearly double the current extent of area-based conservation to 30 per cent of the world surface by 2030 [10], likely to affect the lives and local environments of hundreds of millions of people [11, 12].

Scaling effective and socially just conservation efforts to meet global challenges is an important goal, but little is known about the processes that drive such scaling [13, 14]. Recent research shows that the temporal pattern of scaling in conservation tends to follow the sigmoidal trajectory characteristic of social learning [i.e., peer-to-peer transmission; 13], highlighting the role of neighbour-to-neighbour interactions [15] and key players for information spread [16]. To date, research around scaling conservation has also been largely retrospective [e.g., 13, 14, 15, 17-20]. For example, Romero-de-Diego, Dean [18] interviewed experts to investigate the factors influencing farmers’ adoption of Wildlife Management Units in Mexico. Their study revealed that crucial determinants included the observability of benefits, compatibility with farmers’ needs, accessibility to technical support, and land tenure status. While these studies provide important insights into the factors associated with scaling and the strategies that may support it, there are remaining questions regarding the mechanisms by which initiatives spread and expand, are adapted to new conditions, and persist in different places and over time [16, 21].

Past experiences and trends can also be unreliable guides to how conservation measures may perform in the future, limiting our abilities to pre-empt their impacts in the face of uncertainty and changing conditions. As a result, the science of scaling conservation to meet societal needs risks being reactive rather than proactive [22]. Instead, predictive conservation science approaches can help ensure conservation initiatives are implemented and evaluated in ways that anticipate ever-changing conditions in socio-ecological systems [23]. Predictive research spans a wide range of quantitative and qualitative methods, from scenario-based interviews [e.g., 24, 25, 26] to mechanistic models of complete systems [e.g., 27].

One particularly promising method for predicting the scaling of conservation is agent-based modelling. Scaling is an emergent outcome of smaller, interconnected social and environmental processes that collectively shape the overall patterns of expansion, adaptation, and persistence [28, 29]. In agent-based models, the agents (e.g., individuals or communities) make independent decisions based on predefined rules, iteratively responding to the logical consequences of their past decisions, interactions with other agents, and their environment. Hence, agent-based models are well-suited for exploring how underlying conditions and interactions among individual components affect complex emergent outcomes [30]. This approach allows researchers to delineate their hypotheses about a system’s important underlying mechanisms, test their logic, and simulate alternative scenarios [30].

Agent-based model predictions can be compared for consistency with observed data, such as the times and places conservation initiatives are adopted [31]. Specifically, one can use statistical inference to identify the modelled individual-level processes (e.g., decision rules and parameter values) that match empirical data and thus gain insights into the processes that shaped the observed scaling pattern. Then, with a model that adequately reproduces observed data and thus captures the mechanisms of interest, researchers can forecast future scenarios. This method also allows for predictions across all possible values for unobserved or future data, thus producing estimates of uncertainty based on causal mechanisms rather than canonical measures of variance. This is important, as small alterations in social-ecological conditions can lead to unexpected, disproportionately large changes in outcomes [i.e., they can exhibit chaotic behaviour; 32]. Therefore, agent-based models provide a framework for empirically validating alternative hypotheses about the drivers of scaling conservation initiatives, quantifying uncertainty given these underlying processes, and making informed predictions.

Projections of the spatial and temporal patterns of conservation scaling and associated uncertainties could be useful to conservation practitioners and policymakers in several ways. First, projections could help decision-makers select the most scalable measures from a portfolio of options. These scaling projections could be combined with impact estimates to forecast the cumulative effect of selected conservation measures [33]. These projections may also be useful when communicating with external audiences, such as providing defensible estimates of total adopters when seeking funding. Second, these projections could help understand where and when the ecological and social benefits and costs may likely occur in space and time. For example, when in a project’s timeframe are the highest adoption rates expected, and in what locations? Third, agent-based models are particularly useful when engaging with decision-makers, as they can demonstrate the potential consequences of different policy and management decisions [34]. For instance, decision-makers might explore plausible consequences of changing the design features of initiatives to make them attractive to potential adopters. Finally, such models could be integrated into monitoring, evaluation, and learning systems to track progress and course-correct to scaling targets [35]. For instance, comparing actual to predicted levels of adoption at a given project phase can guide practitioners on when to intervene, such as by adjusting adoption incentives to stay on course with scaling goals.

In this paper, we use data on village engagement with an initiative involving the adoption of Locally Managed Marine Areas (LMMAs) in Fiji to demonstrate how agent-based modelling can be linked with empirical observations to produce reliable inferences and predictions about the scaling mechanisms of conservation actions. LMMAs are areas of nearshore waters and their associated coastal and marine resources that are largely or wholly managed by local coastal communities with support from partner organizations [36]. LMMAs are supported by NGOs, a university and a Secretariat that coordinates village requests for support. The villages and their partners form the Fijian Locally Managed Marine Area (FLMMA) network, which aims to improve the adaptive management of marine resources to benefit livelihood and ecological resources [37]. Villages voluntarily choose to join the FLMMA network and to do so, establish LMMAs in their customary fishing grounds. Specifically, we aim to create an agent-based model that adequately reproduces the spatiotemporal patterns of villages’ engagement with the FLMMA network. Furthermore, we use the important identified processes to forecast future adoption trajectories under different programmatic decisions. We use data on engagement with FLMMAs instead of information on the adoption of LMMAs because the latter is not available for all villages in Fiji.

Using inference techniques to compare predictions from agent-based models with empirical data and subsequently applying these inferences for informed decision-making is nascent in conservation science [e.g., 38]. Thus, we structure the methods section of this manuscript as a step-by-step guide for conducting such analyses. We demonstrate the utility of this approach by providing measures of out-of-sample predictive accuracy and comparing our inference results to previously published work. Specifically, we compared our results against those of Jagadish, Freni-Sterrantino [15], who employed a multilevel regression approach with the same data on villages’ engagement with the FLMMA network. In our discussion section, we interpret our findings, address limitations in our analysis, and explore the implications of this case study and agent-based modelling analysis framework for conservation policy and practice.

## Methods

This paper uses a Bayesian parameter inference approach to arrive at a causal description of why and how engagement with the FLMMA network spreads between Fijian coastal communities. To do this, we construct an agent-based model depicting the dynamics and mechanisms that govern the adoption process. Agent-based models contain a set of free parameters, and the inference task is to learn the ranges of parameter values that ensure that the model predictions reproduce empirical observations. Since the different parameters are associated with different mechanisms (e.g., whether information flows between villages), the parameter intervals that accurately replicate the observations shed light on the mechanisms driving adoption in a system.

In this methods section, we go through the steps to build and apply an agent-based model of conservation dynamics within a Bayesian framework, as follows:

**Step 1: Building the conceptual model that describes the adoption process:** The first step is building a conceptual model of the dynamics that drive the outcome of interest. Below, we present a conceptual model developed in collaboration with Fijian practitioners, scholars, and rights-holders.

**Step 2: Translating the conceptual model into an agent-based model:** The second step is translating the conceptual model into a mathematical model in the form of a computer simulation that predicts adoption patterns. In this paper, we create two agent-based models to simulate different aspects of individual villages’ behaviour: a stochastic agent-based model and a deterministic agent-based model (described below).

**Step 3: Defining the fitting criteria:** The third step is to define how to compare the model predictions to real-world data from our case study (i.e., our empirical observations). In other words, we decide what aspect of the data the model should replicate and how to quantify the model fit. In Bayesian statistics, this comparison is performed by computing the likelihood: the probability of obtaining the observed data given the model parameters.

**Step 4: The inference framework:** Having constructed our agent-based model (step 2) and defined the fitting criteria (step 3), we proceed to the Bayesian parameter inference (i.e., we find the parameter values that adequately reproduce empirical observations). Specifically, we obtain the joint posterior probability distribution, which allows us to quantify the credible intervals of the parameters. To accomplish this task, we run the model with different combinations of parameter values and evaluate how well each parameter combination performs. Because the number of parameters in our model is large, a brute-force trial-and-error procedure across all parameters is computationally insurmountable. To overcome this issue, we rely on Monte Carlo sampling algorithms that explore different parameter values while autonomously avoiding less promising parameter combinations. In addition, we limit the parameter search to realistic parameter ranges given the phenomenon of interest. Within Bayesian statistics, such *a priori* parameter constraints are called priors.

In addition to providing parameter estimates, the obtained causal model allows for robust out-of-sample predictions. This approach allows us to explore counterfactual scenarios and make predictions about the future.

### Step 1: Building the conceptual model that describes the adoption process

We developed the conceptual model describing the adoption process based on the existing literature and iterative discussions with representatives from the FLMMA network. If, during discussions, a specific characteristic was identified as potentially influencing the adoption and spread of the initiative, and we could identify a variable to represent that process, it was included in the model.

Engagement with the FLMMA network is a voluntary process that must be initiated by community representatives. A ‘community’, in this case, is often represented as the village and thus assumed to be the village in this study. We assumed that villagers must first learn about the FLMMA network before engaging (A circled in blue in Figure 1). Villagers can learn about the FLMMA network through a range of channels, including media (B circled in blue in Figure 1), interactions with neighbours (including with villages that are physical neighbours, share schools, churches, hospitals, and customary fishing grounds (or *qoliqoli*), or attend the same district meetings; C circled in blue in Figure 1), or via supporting organisations (D circled in blue in Figure 1).

**Figure 1:**
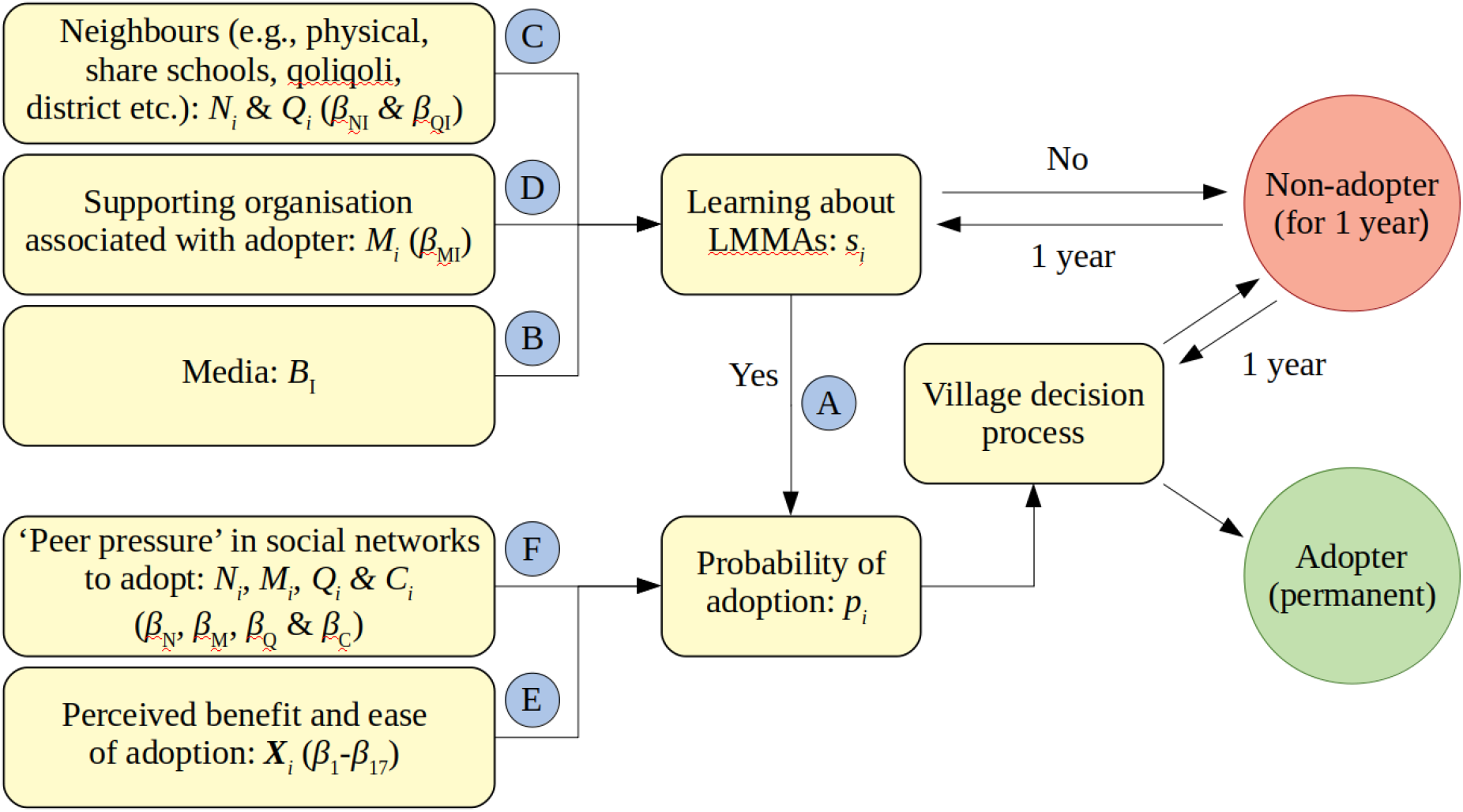
Flowchart summarising our conceptual model for adoption. The variables and parameters listed in the different boxes refer to the equations listed in Step 2. Note that the parameters β_NI_, β_MI_, β_QI_, *B*_I_, and *s*_i_ only appear in the deterministic version (Step 2.2), while the same mechanisms are included but modelled in a different manner in the stochastic version (Step 2.1). The letters in the blue circles are used in the description of the flowchart in the main text.

Once village members have learned about the FLMMA network and are interested in adopting LMMA, they will approach FLMMA representatives through an official letter of interest endorsed by the village chief. Various factors are expected to influence a village’s perceived benefits and ease of adopting an LMMA and engaging with the FLMMA network. These include endogenous characteristics, such as the perceived extent and diversity of benefits of adopting, the degree to which LMMAs are compatible with local needs, perceptions of political empowerment, and the availability of technical support [E in Figure 1; e.g., 15, 39]. These factors also include exogenous characteristics, such as whether their neighbours or other connections (e.g. vilages that share schools or *qoliqoli*) have previously adopted, thus exerting “peer pressure” to adopt [F circled in blue in Figure 1; 15, 39].

Next, the village creates a management plan through a series of workshops [40]. Support organisations, including NGOs, the Ministry of Fisheries, and a university partner, facilitate (influence E circled in blue in Figure 1) initial engagement with the FLMMA network and help the community identify the issues and solutions they wish to address. However, their presence is not mandated for engagement [40]. Finally, the village chief, in consultation with the community, will decide whether to engage with the FLMMA network and thus adopt an LMMA (i.e., village decision process in Figure 1).

Multiple villages can also engage with the FLMMA network as a group (impacting C, D, and F circled in blue in Figure 1); thus, one village can influence the decisions of other villages to engage. For example, where multiple villages share a communal fishing area, supporting organisations will consult with all villages. Often, the village with the highest status within traditional governance systems (the “chiefly village”) will lead the engagement. Additionally, many communities can work together to establish a Resource Management or ‘*yaubula’* committee to oversee the development and implementation of their LMMA. These committees can represent multiple villages at a district or provincial level; in these cases, each village has a representative on the committee.

Supporting organisations also have a process to decide whether they will support a community or not. This process includes assessing a community’s social and environmental conditions and whether establishing an LMMA is necessary or sufficient to address ongoing issues. In practice, few community applications to join the FLMMA network are rejected. Thus, in our conceptual model, we assume that if a village decides to participate, it will be admitted into the FLMMA network. While we know there are many cases of LMMA abandonment, data on these are not available. Thus, our model assumes that the decision to adopt an LMMA is permanent. Importantly, given our lack of data, our model does not account for the differences in engagement with supporting organisations or the strategies they use. We do not consider the impact of external processes (e.g., the amount of money being invested by different support organisations) and partners as these data were not available; however, we know this is likely to be significant [41].

### Step 2: Translating the conceptual model into an agent-based model

The features described in the previous section are expected to influence the uptake of conservation initiatives as a function of time and geographical location. Thus, the modelling framework described below aims to capture both the temporal and spatial adoption dynamics. We present two models: a deterministic and a stochastic version. The deterministic model tracks the probability of adoption for each village as a function of time. In contrast, the villages in the stochastic model make a binary decision of whether to adopt at each timestep. The deterministic model can be employed for Bayesian inference around the influence of each variable [42], suggesting factors that most influence the pattern of scaling. In contrast, the stochastic model can be used as a generative model that produces synthetic data, allowing users to explore plausible future management scenarios. In our analysis, we employ both agent-based models, referring to the stochastic model as sABM and the deterministic model as dABM.

#### Step 2.1: Stochastic agent-based model (sABM)

We start by describing the stochastic agent-based model, where we consider each village as an independent agent that can decide to adopt the conservation initiatives at each timestep. We use a timestep of one year. While villages are assumed to stick to their decision if they adopted the initiative at a previous timestep, villages who do not adopt reevaluate their decision every timestep. The probability (*p_i_*) that the *i*th agent will adopt the measure within a given timestep depends on the perceived benefits and ease of adopting (***X****_i_*) (E in Figure 1) and the number of various types of ‘neighbours’ that have previously adopted the initiative (F in Figure 1). We assume that as more ‘neighbours’ adopt, the pressure on the village to adopt increases. More specifically, we assume that (eq. 1)

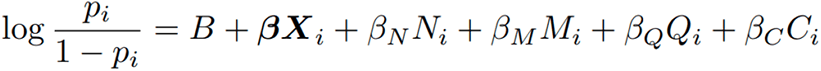

Here, **β** is a vector containing the free parameters associated with each variable influencing the perceived benefits and ease of adopting (***X****_i_*). *N_i_* is the number of direct neighbours already adopting the initiative, and β_N_ denotes the associated free parameter. Thus, we assume that the greater the number of immediate neighbours who engage with the FLMMA network, the more likely it becomes for a village to engage themselves. We assume villages are neighbours (*N_i_*) if they lie within a distance shorter than 1.5 times the distance to the closest geographical neighbour or if they share a school, hospital or church within 5 km.

In eq. (1), we also assume that villages will be more likely to adopt the initiative if they have an existing link with a supporting organisation through which they encounter other adopters. However, we assume the impact of connections to adopter villages through the supporting organisation to be different from that of direct neighbours. Thus, *M_i_* denotes the number of adopters associated with the same supporting organisation as the *i*th village, and β_M_ denotes the associated free parameter. A direct neighbour might also be a member of the same supporting organisation, impacting the village’s decision through both *N_i_* and *M_i_*. Meanwhile, *Q_i_* denotes the number of villages that share the same customary fishing ground, while *C_i_* indicates whether the chiefly village is an adopter, as these links are likely to influence the adoption of other related villages. While the different social networks (associated with *N_i_*, *M_i_*, *Q_i_* and *C_i_*) are not identical, they are correlated, and we note that this might introduce ambiguities for the interpretation of the fit as they are not straightforward to disentangle. Finally, *B* is a base-rate probability for adoption.

Before villages can adopt the conservation initiative, they must first learn about it. In our model, some villages will learn about the FLMMA network from the onset, for instance, if a support organisation is present (D in Figure 1). Other villages randomly learn about the FLMMA network with a fixed base-rate probability (e.g., representing information transmitted over the phone or TV; B in Figure 1), and others are assumed to learn about the FLMMA network if their connections (either *N_i_*, *M_i_* or *Q_i_*) (C in Figure 1) have engaged. Thus, upon adoption, the villages are assumed to communicate their decision to their network.

Note that *N_i_, M_i_* and *Q_i_* in eq. (1) include villages that have adopted the measure during the same timestep (i.e., the same year). Moreover, it is reasonable to assume that the information about the adoption by villages will spread across their social network upon adoption (i.e., within the same year). Thus, to account for the impact of the village on other villages *within* the same timestep, we loop through the villages in random order at each timestep to minimise biases.

#### Step 2.2: Deterministic agent-based model (dABM)

Our aim in constructing this agent-based model is to compare our model to real-world data and to use this comparison to infer model parameters and gain insights into the dynamics that underlie the system. However, performing Bayesian inference using stochastic agent-based models is notoriously hard (e.g., when the agents’ behaviour is governed by random processes, such as the random spread of information discussed above (see also Jørgensen, Ghosh [43] for more details on the topic)).

To circumvent this issue, we created the deterministic version of the agent-based model. We do so by adding a few simplifying assumptions. First, we assume that a village’s decision will only impact other villages’ decisions (through *N_i_*, *M_i_* or *Q_i_* in eq. 1) if they adopt the initiative before the present timestep. In other words, we assume that villages need to see the successful implementation of the initiative over a certain timescale before it weighs into their decision. By introducing this assumption, it is no longer necessary to randomly shuffle the order in which we compute the adoption probabilities of the villages at each timestep. Secondly, for the purpose of Bayesian inference, we only need to reliably recover the probability that a village will adopt as a function of time rather than asking the village to make a binary decision based on this probability. Thus, we record the probability that a village will have been informed and adopted at or before each timestep.

As shown in the results section, the impact of these assumptions on the posterior prediction of the model depends on the model parameter values in question. However, in many of the scenarios presented in the results section, the deterministic model yields a good approximation to its stochastic counterpart.

For the deterministic model, we still use eq. (1) but replace *N_i_*, *M_i_*, *Q_i_* and *C_i_* with the corresponding probabilities (e.g., a neighbouring village that will have adopted with probability 0.1 counts as 0.1 neighbours in *N_i_*). Finally, we must deal with the spread of information since this aspect likewise has a stochastic component in the original model. We note that the probability of a village having learned about the measure *and* adopted it equals the probability of adoption, assuming that the village has learned about the measure (i.e. eq. 1) times the probability of learning. As discussed above, the probability of learning (hereafter *s_i_*) at the present timestep depends on the network of the villages and includes a fixed base-rate probability. Moreover, assuming that learning is binary, we can assume that (eq. 2)

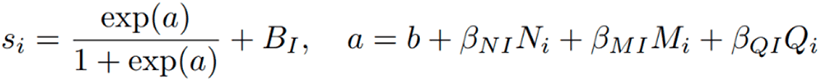

where *B_I_*, β*_NI,_* β_MI_ and β_QI_ are free parameters, and *b* is a fixed negative scalar that sets the scale for the free parameters and ensures that *a* is negligible when *N_i_*, *M_i_* and *Q_i_* are zero. Instead of a stochastic model for the spread of information, we thus now have an equation that assigns a probability to each village for learning during the present timestep. The probability (*S*_i_) of learning at or before the present timestep (*t*) is then (eq. 3)

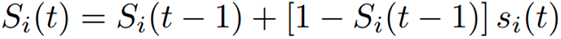

where *s_i_* is the probability computed by eq. (2). Finally, the probability of having adopted by or during the present timestep for the *i*th village is then (eq. 4)

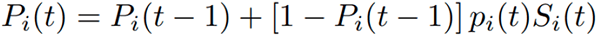

where *p_i_* is given by eq. (1) and *S_i_* stems from eq. (3).

All model parameters used in the deterministic and stochastic versions of our agent-based model are summarised in Table 1.

**Table 1:**
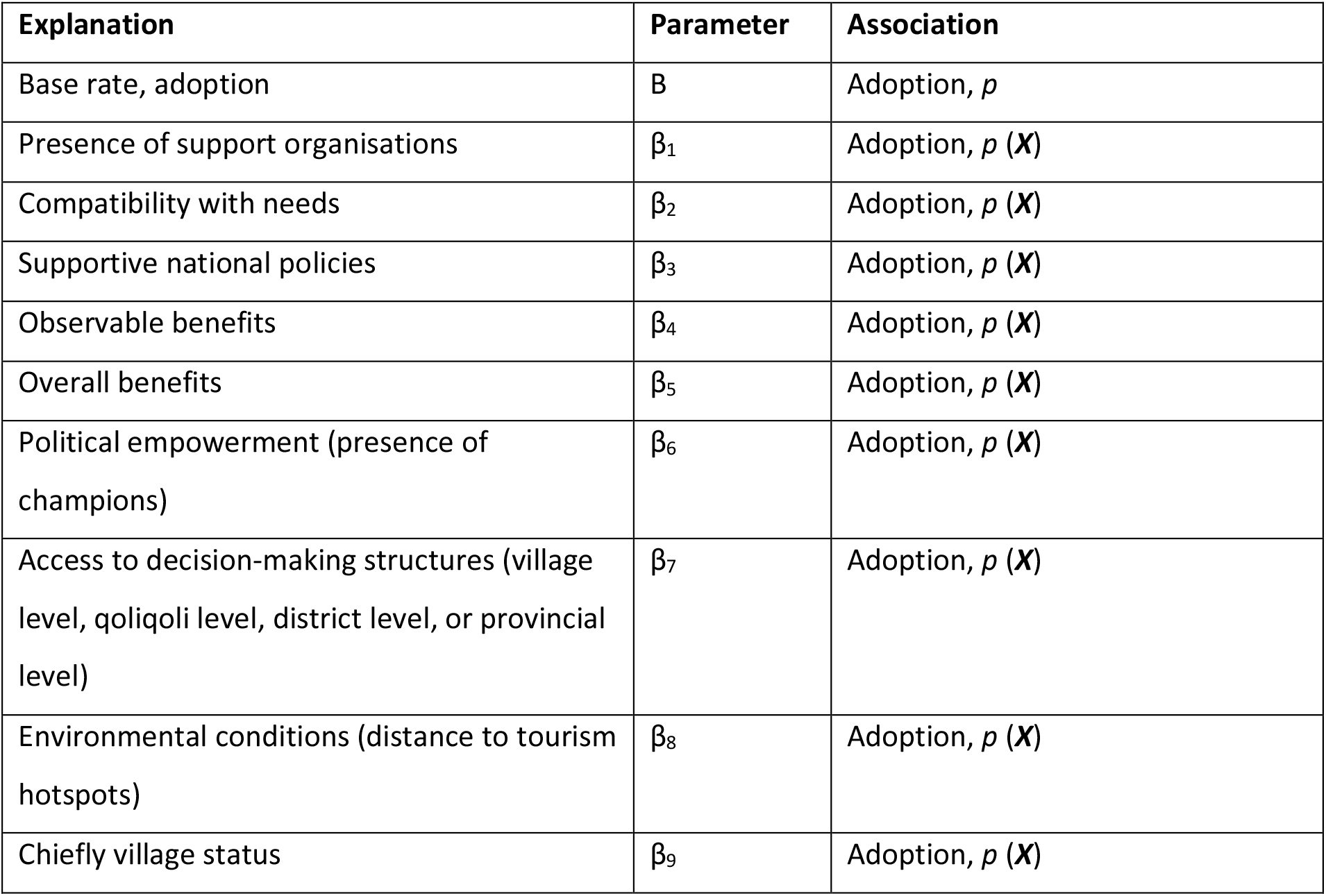
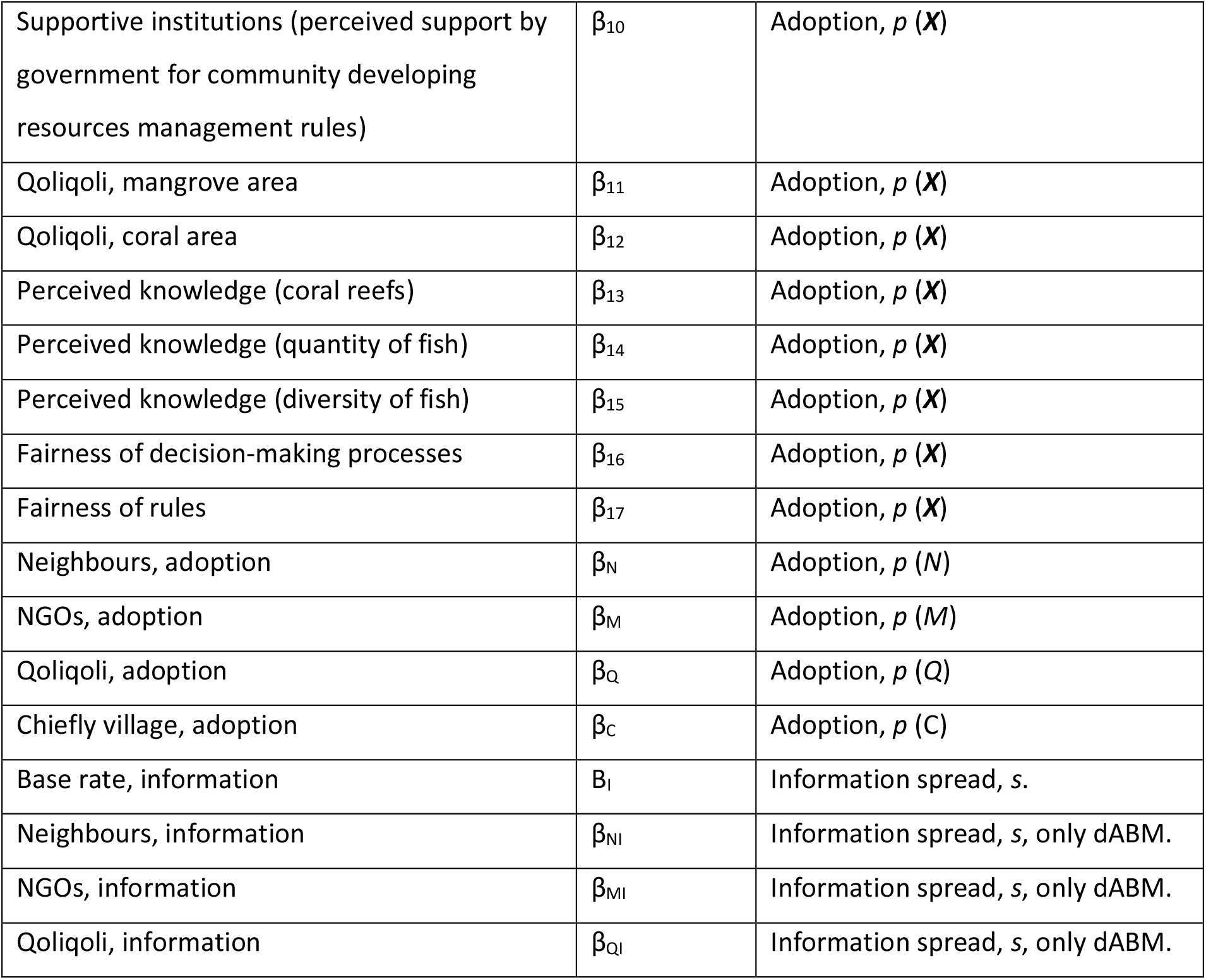
Table of parameters. All information for β_1_-β_17_ is available for the 122 villages in the data set discussed in this paper. We include another data set with 627 villages in Appendix C. For this latter data set, only *B*, β_11_, β_12_, β_N_, β_M_, β_Q_, *B*_I_, β_NI_, β_MI_, and β_MI_ were available. Details of all survey data are provided in Table S1.

### Step 3: Defining the fitting criteria

To compare our model to data, we need to define the likelihood function, i.e., the probability of observing the actual adoption pattern based on the model predictions for a given set of model parameters. In our model, each village faces a binary decision at each timestep, leading to adoption with probability *P_i_*. When comparing the model predictions to the data on an individual basis (i.e., when including the spatial information), we can compute the joint probability of the observed adoption pattern as the product of the individual adoption events. We thus attribute the probability *P_j_* to *j*th observed adopter and the probability (1-*P_k_*) to the *k*th observed non-adopter. By doing so, we track the exact timeline of events and make full use of the spatial information. We follow this procedure for several fits presented in the results section below. Thus, the logarithm of the likelihood (log*L*) is given by (eq. 5)

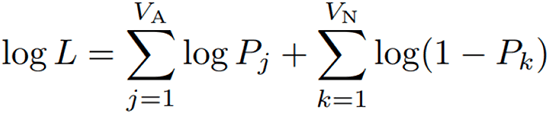

where we sum over the *V*_A_ observed adopters and the *V*_N_ observed non-adopters.

To understand the value of our data to make predictions at the level of individual villages, we repeat the analysis using alternative fitting criteria. Hence, imagine that we did not actually know the identity of adopters but merely had access to the total number of adopters as a function of time. We know the information stems from a set of Bernoulli processes with individual probabilities *p_i_*. However, we do not know which Bernoulli process leads to which adoption event. If all villages adopt at the same rate, the probability of observing a given number of adoptions as a function of time could be computed using a binomial distribution. We might thus make the simplifying assumption of a constant adoption rate equal to the mean adoption rate at the time point, i.e., given that the likelihood is intractable, we introduce a surrogate likelihood on the form (eq. 6)

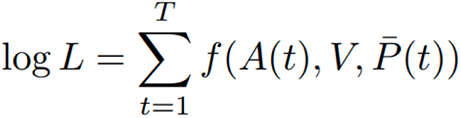

where *A*(*t*) denotes the number of adopters at time *t*, *V* denotes the total number of villages in the data set, and *p̅*(*t*) is the average value of the adoption probability at time *t*, and the function *f* denotes the logarithm of the probability mass function for a binomial process. We sum over all *T* time points. The task is then to maximise the probability (eq. 6) of producing the observed adoption pattern by choosing suitable model parameters. We present results following this approach for a larger data set for which the features (***X***) are unavailable. However, this approach does not make full use of spatial information. The model is thus not required to reproduce the heterogeneity of the villages in the data set but merely must recover the accurate number of adopters using as few parameters as possible.

In this paper, we apply our framework to the data set discussed by Jagadish, Freni-Sterrantino [15], representing 146 villages in Fiji located within 1 km of the coastline for which Ministry of Lands and Resources spatial information were available. From this data set, we selected the 122 villages for which all variables associated with the parameters in Table 1 were available. These villages are on the islands of Viti Levu, Vanua Levu, Kadavu and in the Lomaiviti group. These island groups were selected because of accessibility and budget for data collection. Villages are identified as part of the FLMMA network based on the list of villages in the FLMMA 2020 database. Non-FLMMA network villages were selected based on one-to-one propensity matching without replacement, so they resembled FLMMA network villages regarding covariates, which may influence adoption and impacts [for additional information, see 44]. Our data set also included spatial data on coral reef cover [45] and the 2016 mangrove cover data from the Ministry of Forestry within *qoliqoli* (traditional fishing ground) boundaries.

Data were collected between October 2019 and March 2020 through interviews with groups of key informants representing village leaders. Participants were selected through consultation with village chiefs and other village leaders and in consultation with additional community members during a traditional *sevusevu* ceremony. Key informants represented three types of leaders: *vanua* (land people, custom), *lotu* (spiritual) and *matanitu* (government). Surveys were piloted before they were deployed. Interviews lasted an average of 75 minutes. Consent was obtained from the village chief and key informants before interviews were conducted, and ethics permission was obtained (Middlesex Ethical Review Board Application 8030). All data collected and selected for the models were based on the Diffusion of Innovation theory [see 15 for more details] and feedback from local experts on how communities learned about the FLMMA network.

### Step 4: The inference framework

To perform the inference task, we compare the model predictions to data for different combinations of parameter values. The technical details of how this is done in practice are summarised below.

We use the Markov chain Monte Carlo (MCMC) ensemble sampler presented by Foreman-Mackey, Hogg [46] and the Gibbs sampling algorithm SLInG (Sparse Likelihood-free Inference using Gibbs [47]). The latter imposes sparsity-inducing priors; the algorithm autonomously applies Occam’s razor, searching for suitable fits with a low number of non-zero parameter values. For the MCMC ensemble sampler, we impose uniform priors, limiting the region of interest to values between 0 and 40 for all parameters except for B*_I_* that can take values between 0 and 1 and B, β_7_, β_9_, β_11_, and β_12_ that are allowed to take values between –40 and 40.

The MCMC ensemble sampler explores the space using several walkers, i.e., it explores different regions of the parameter space simultaneously, thus reliably capturing multi-modal distributions (see Figure S6). Thus, we are not limited to exploring the parameter values that lead to the maximum likelihood. Rather, the obtained joint posterior distribution maps the distribution of parameter value combinations that are consistent with the empirical observations and the prior. In contrast, SLInG maps the posterior distribution for a sparse model that is consistent with the data. The solution obtained with SLInG is hence not unique and reflects the level of sparsity induced through the prior.

## Results

### Predicting current adoption

In this section, we show that our model successfully reproduces the spatiotemporal patterns of the spread of the FLMMA network across villages in Fiji. Moreover, we estimate the empirical values of the model parameters and their associated uncertainties. We present results obtained from the data set that includes 122 villages for which information on the perceived benefits and ease of adoption (β1-β17) was available.

First, both the deterministic and stochastic models reasonably fit the observed cumulative adoption pattern (Figure 2, cf. the performance metrics in Figures S1 and S2). The deterministic version of the agent-based model yields a good approximation to the behaviour of its stochastic counterpart, with which we get a realistic estimate of the model uncertainties (Figure 2). The uptake of the initiative is slightly higher in the stochastic model, which we attribute to the fact that villages in the stochastic model are affected by other villages that adopt within the same timestep.

**Figure 2:**
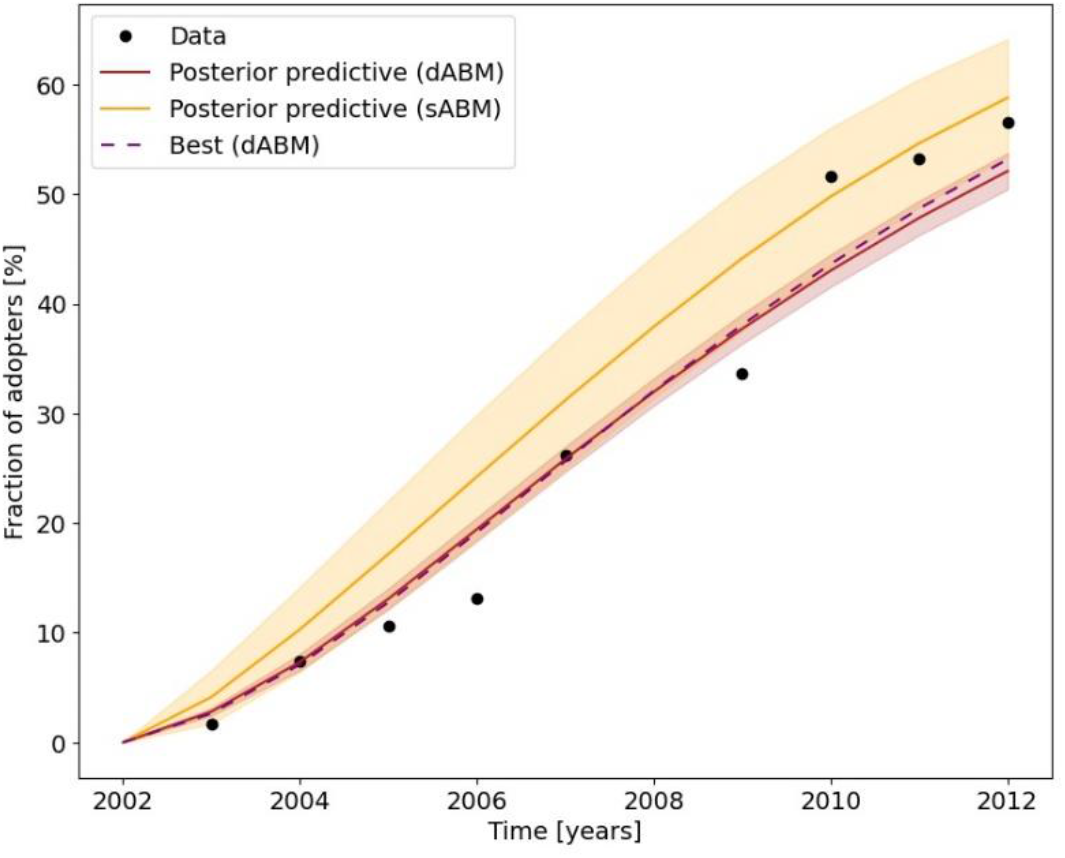
Adoption as a function of time for the fit to the deterministic agent-based model (presented in Figure 3a) and stochastic model. The shaded area signifies 95 per cent credible intervals, while the solid line shows the corresponding median. The dashed line shows predictions by the best parameter set among models in the Markov chains.

We repeated our analysis using SLInG (i.e., imposing sparsity-inducing priors, as this allows us to single out a subset of the parameters that can explain most of the variance in our data. Doing so, we find that only the base rate of adoption (B), the overall benefits (β_5_), the perceived knowledge of fish diversity (β_15_), the network of supporting organisations (β_M_), and the base rate of information spread (B_I_) are needed to explain the data (cf. Fig. 3b); these four parameters are associated with perceived benefits and ease of adoption. Under the assumption that these variables are causally independent and that unobserved confounders do not influence our model, these four variables thus stand out as potential main drivers of adoption. We will address this assumption further in the discussion section.

**Figure 3:**
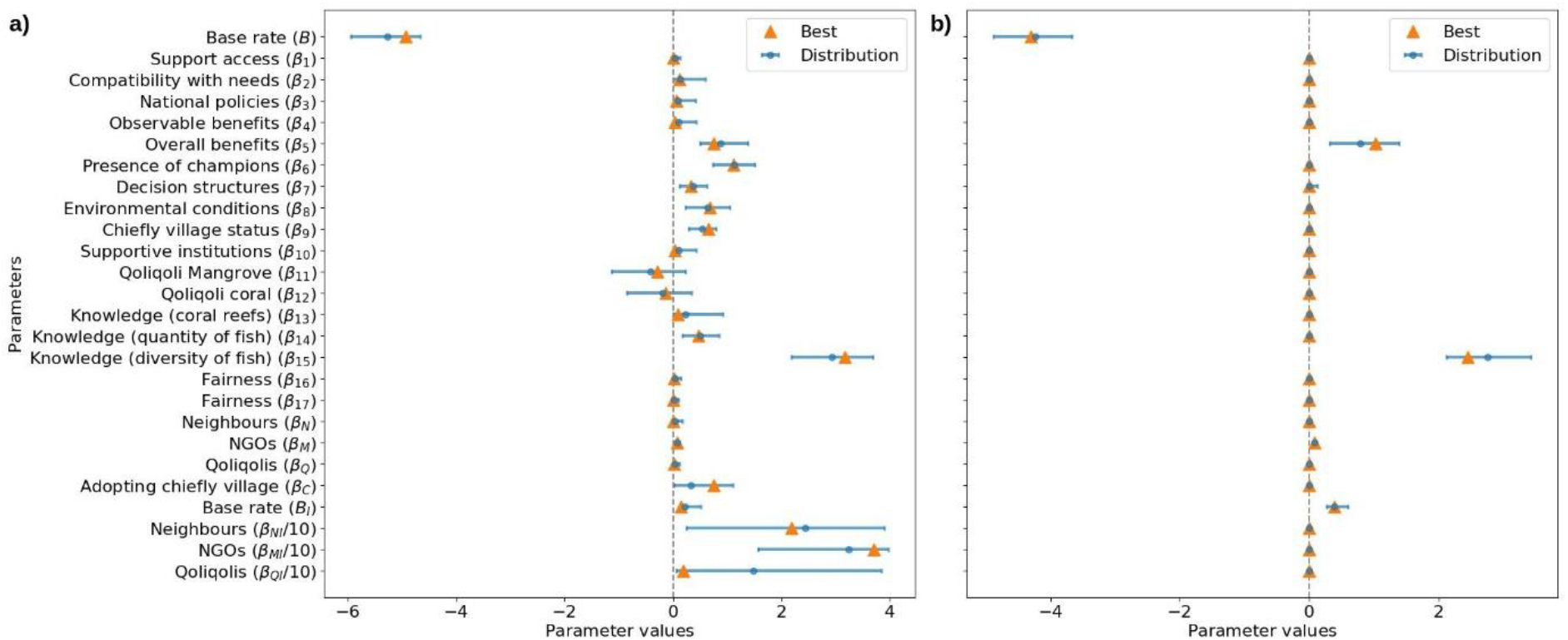
Summary of the posterior distribution of the inferred model parameters. The error bars indicate 95 per cent credible intervals. The circle shows the median, while the triangle represents the best fit. a) The fit was obtained using the MCMC ensemble sampler based on 13 million simulations, excluding the burn-in. b) We repeated the analysis using SLInG, finding that the only non-zero parameter values left (after imposing sparsity priors on all parameters except the base rate of adoption (B) and base rate of information spread (B_I_)) were the base rate of adoption (B), overall benefits (β_5_), perceived knowledge on diversity (β_15_) of fish, the impact of network of the supporting organisations (β_M_), and the base rate of information spread B_I_.

When we repeat the analysis but fit only the total number of adopters as a function of time (i.e., neglecting the spatial information), the model parameters associated with the perceived benefits and ease of adoption are no longer required to reproduce the observed data. This change in parameter importance and loss in constraining power is because the individual villages have become largely interchangeable for the purpose of the fit. We also note that the approximations behind the deterministic version of the agent-based model do not yield a good approximation to our stochastic model in this case. However, the model fits the total number of adopters as a function of time with high accuracy.

Repeating the fit to the cumulative adoption as a function of time using sparsity-inducing priors, we find that only the base rates for adoption (*B*) and information spread (*B*_I_) are needed to explain the data (cf. Figure S3). However, the obtained parameter values do not predict the spatial distributions of adopters as well as the fit presented in Figures 2 and 3 (cf. Figure S1 and S2).

In addition to the data set presented above, we analysed a larger data set containing 627 villages, obtaining results consistent with those discussed above. These results are summarised in Appendix B and C.

### Predicting future adoption

We can project the model into the future by using the parameter values obtained from fitting our agent-based models to data. We use this projection to investigate the impact of potential interventions. Doing so allows us to guide policies that can increase future uptake of the conservation initiative. For instance, the model presented in Figure 3a attributes a significant effect to the presence of champions (β_6_). To quantify the impact of changes associated with the presence of champions on future adoption, we can simulate the consequences of these policies using contrafactual values of the survey responses associated with this parameter. We will thus either assume that all champions are withdrawn or that the presence of champions is sharply increased. The results of such an investigation are shown in Figure 4. As can be seen from the figure, future uptake is expected to be dampened if villagers did not recognise champions for resource management within their community. Thus, while the median fraction of adopters is forecast to be 86 per cent in 2030 when champions for resource management are present, a withdrawal of all champions would lead to a median uptake of only 75 per cent by the same year.

**Figure 4:**
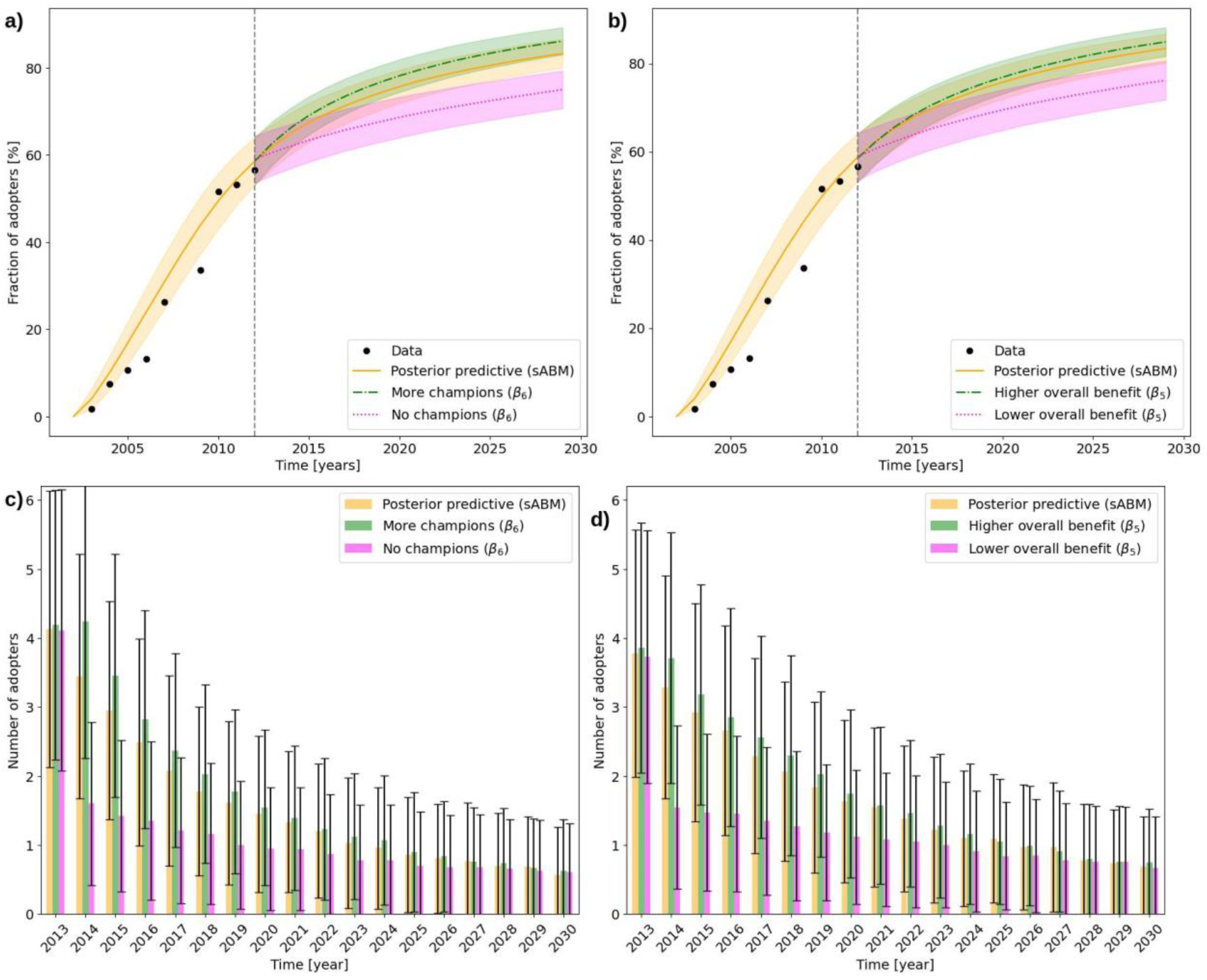
Different future scenarios based on the fit presented in Figure 3a. a) The impact of the presence of champions on future adoption. Here, we either assume that all champions are withdrawn or that the number of champions increases. b) The impact of the perceived overall benefit on future adoption. We either assume that all villages agree that the overall benefit corresponds to the highest overall benefit reported within the sample or to the lowest value within the sample. c) Bar plot showing the number of new adopters as a function of time when changing the number of champions. d) Like c but for the different scenarios associated with a change in the perceived overall benefits.

As another hypothetical future scenario, we investigate the changes in the overall benefit by using counterfactual values for each village (β_5_). We thus either assume that all villages agree that the overall benefit corresponds to the highest assessment of the overall benefit across the villages or to the lowest assessment in the sample. The results are shown in Figure 4. As can be seen from the figure, a reduction in the perceived overall benefits would severely dampen future uptake, leading to a median uptake of only 76 per cent by 2030. In contrast, with high perceived overall benefits, the median fraction of adopters is projected to be 85 per cent by the same year.

Finally, we investigate the impact of three scenarios reflecting different investments in media about LMMAs (our base rate information spread, *B*_I_). In our model (cf. Figure 3), information spread is dominated by the information spread through media, i.e. the base rate (*B*_I_), and information spread through supporting organisations (β_M_). First, we investigate a scenario in which there were no media present from the onset (Figure 5, dash-dotted line). This leads to a drop in uptake across the entire time series. In the second scenario, we assume that there is no further media coverage of LMMAs after 2012 (Figure 5, dotted line). This does not impact the uptake, as all villages are informed about LMMAs by this time. Finally, we look at a scenario where information can only spread through the media (Figure 5, dashed line). Again, this impacts the total uptake across the entire time series.

**Figure 5:**
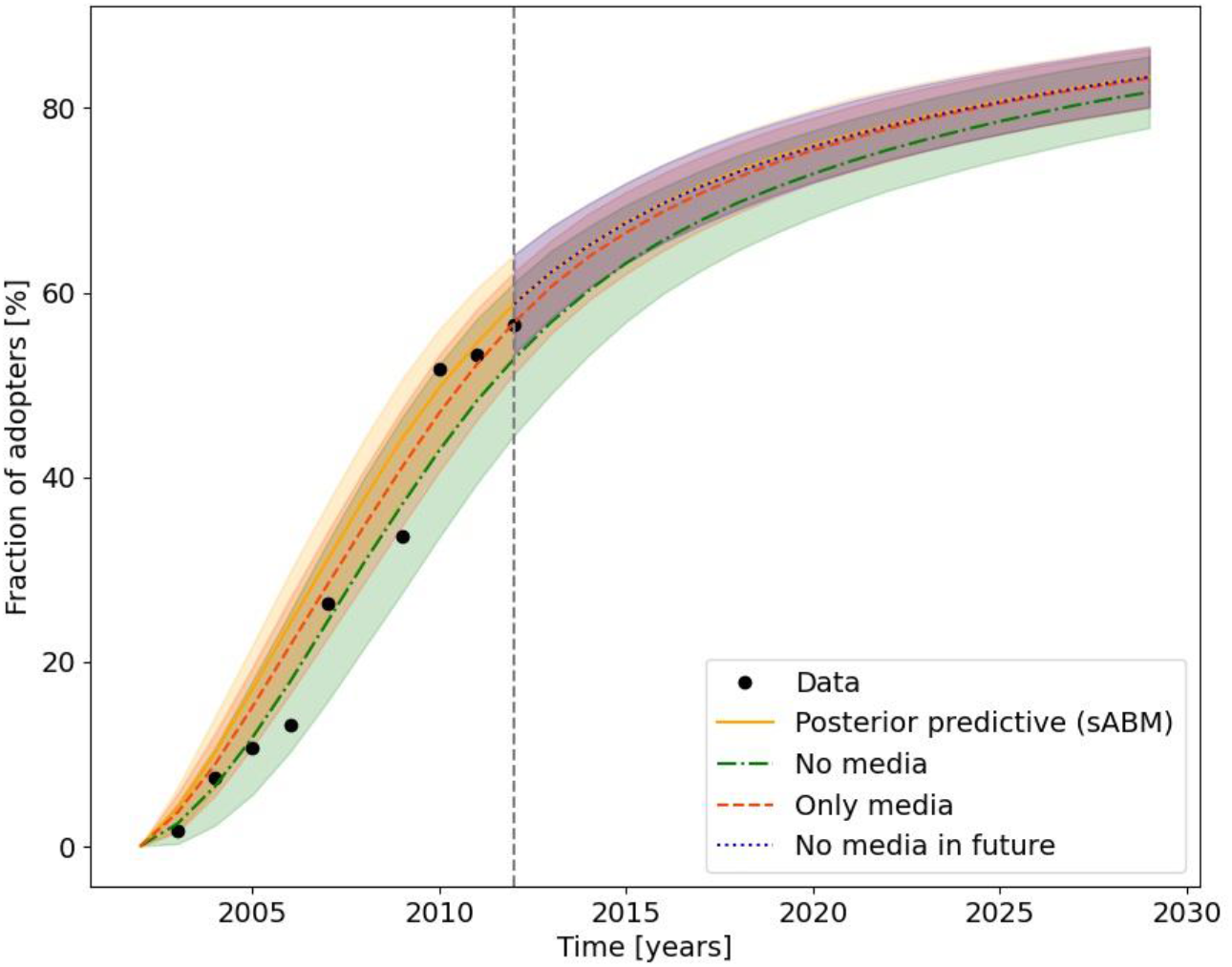
Different future scenarios for different investments in general media based on the fit presented in Figure 3a. We illustrate the impact of three counterfactual scenarios: no media present from the onset (dash-dotted line), no further media coverage of LMMAs after 2012 (dotted line), and information can only spreads through the media (dashed line).

## Discussion

Efforts to scale conservation action to reverse global nature loss and tackle climate change while supporting human well-being have been inadequate [1, 2]. While an emerging body of research studies how conservation initiatives go to scale [e.g., 13, 14, 15, 17-20], there has been little investigation of how to scale conservation actions under future scenarios and uncertainty. We provide one of the first studies using predictive conservation science methods (agent-based models) integrated with a robust theoretical framework (Diffusion of Innovations theory), existing literature and local expertise to forecast the adoption and spread of conservation actions. A key benefit of causal models, such as the agent-based modelling approach presented here, is the ability to produce within-sample predictions regarding counterfactual scenarios and out-of-sample forecasts about the likely pattern of future adoption across space and time (cf. Figure 4).

We show that this modelling framework can accurately reproduce the observed spatiotemporal adoption pattern for engagement with the FLMMA network from 2000 to 2018. Using Bayesian inference techniques, we identify which factors (and the associated parameter value intervals) combine to accurately mimic the observed spatiotemporal spread of the FLMMA network. Moreover, we demonstrate how our modelling approach can help forecast plausible consequences of different conservation policies and practices on future scaling trajectories.

Our results have several key implications. First, accurately forecasting how, where, and when community conservation practices will be adopted may be challenging without detailed and specific knowledge about the communities involved. For example, we needed information on how villages are connected to each other and the characteristics of the villages (e.g., they perceive overall benefits associated with LMMAs, and perceive greater knowledge of fish diversity) to accurately reproduce the spatial patterns of spread. Conversely, the temporal trend in cumulative adoption can be produced by many different spatial processes; i.e., the time series of adoption are equifinal [48]. Therefore, in this case, simply tracking how many villages adopt something over time does not tell us which villages will likely engage in the future. In contrast, including spatially explicit information helps identify the mechanisms driving adoption. Importantly, the base rate of information spread (representing information from, e.g., radio, TV or a university student coming home and talking about the FLMMA network) played an important role in defining the temporal rate of spread, indicating its utility as a mechanism to encourage adoption. Our model could be further improved by capturing information on the actions and strategies of FLMMA network partners. At the moment, all support organisations impact the spread of engagement with the FLMMA network in our model homogenously by facilitating the adoption process or spreading information, but their influence (e.g., how they work, funding, priorities) varies widely, and we do not account for how this might impact the spread. For example, there are stark differences in funding for LMMA development across sites in Fiji [41].

More broadly, key aspects of our model depend on developing a locally appropriate conceptual framework informed by expert knowledge, with implications for the generalizability of our model. We believe the overall structure of our model – namely, the measures of the probability of adoption and the probability of learning about the initiative – are likely to be applicable across contexts (though further research is needed to test this generalizability). However, the specific variables and their values – such as those contributing to perceived benefit and ease of adoption – will vary between contexts. For example, the strength of land tenure is often an important determinant of farmers’ willingness to invest in nature-friendly practices [e.g., 18, 49, 50], while the importance of chiefly village status is specific to our case study (cf. Figure 3a). Villages with chiefs of higher status may be more likely to adopt because those chiefs are more attuned to national debates around fisheries or may be targeted by the FLMMA network. Consequently, models for accurately forecasting spatiotemporal adoption patterns may require detailed empirical data, ongoing collaboration with local experts, a focus on appropriate spatial and temporal scales, and the flexibility to adapt models as systems evolve.

Overcoming these constraints, our models can provide valuable insights, such as among which groups and locations the social and ecological benefits and cost of conservation actions are likely to occur. For example, with and without using sparsity priors, our deterministic model suggested that communities perceiving greater benefits from adopting LMMAs, and with greater perceived knowledge of fish diversity, were more likely to adopt. Consequently, NGOs wishing to facilitate the scaling of the FLMMA network may prioritise these locations first.

Second, well-designed models may help forecast how changes in one characteristic of the system may impact the spread of conservation measures, such as the FLMMA network, into the future. Such insights could be integrated into monitoring, evaluation, and learning systems to track progress and course-correct to scaling targets. For example, based on our inference results, we explored the consequences of altering the number of local champions on projection adoption trends. If an NGO finds itself off-track to meet scaling targets, it might look for ways to encourage and support local champions. In Kadavu in Fiji, this is done by improving environmental awareness of the specific group (e.g., leadership, women) within the community and the community as a whole and by recruiting community members as part of the core team [51]. Of course, the extent to which local champions can be developed depends on the willingness of local people, so it is not necessarily something an NGO can control. More broadly, developing forecasts for plausible adoption trends could help decision-makers select the most scalable measures from a portfolio of options. For example, repeating our approach with alternative fisheries interventions – such as a pay-to-release by-catch intervention [26] – could help anticipate which measure is more likely to be widely taken up. Such projects could be combined with evidence from impact evaluations, such as the national scale evaluation of the FLMMA network by O’Garra, Mangubhai [44], to calculate the relative cumulative impact of different measures. Furthermore, such forecasts are useful when engaging with decision-makers [34].

In a companion study, Jagadish, Freni-Sterrantino [15] analysed a data set including 146 villages using multilevel regression, including a subset of the parameters (β1-β10) in our analysis. Their results are overall consistent with those shown in Figure 3a. In agreement with Jagadish, Freni-Sterrantino [15], we find the perceived benefits of adopting and links to supporting organisations were significant predictor of adoption. This agreement is reassuring, showing that the list of identified significant predictors is consistent with a broader range of modelling choices.

### Study considerations

In this paper, we have introduced two simplifying assumptions, both because of a lack of data. First, the perceived benefits and ease of adoption for each village are assumed to be static properties. However, these will likely change over time due to changing social and environmental contexts, including as the result of the scaling of LMMAs [33]. Data on the time evolution of the perceived benefits are, however, not available. Second, we assume that a village that adopts an LMMA via the FLMMA network does not later abandon it, although LMMAs in Fiji, along with other conservation interventions across the globe, show non-negligible rates of abandonment, especially as a result of environmental or economic shocks. However, abandonment and engagement might take many different forms (e.g., partial abandonment), and data were only available on the initial uptake [53]. Future work might integrate both changes in perceived benefits and abandonment dynamics into our model. Of particular interest would be a model that captures these processes as driven by changes in local environments or climates. Such work might valuably inform predictions of the probability of reaching both local and global conservation uptake targets in the face of increasing climate variability and environmental degradation.

Mathematical models, like the agent-based models presented here, assume that researchers know and can accurately depict the key underlying processes of the system [52]. In complex social-ecological systems, this can be challenging. Crucially, the accuracy of our forecasts is conditional on the assumption that the factors we identified are causally related to adoption and that unobserved confounding variables do not influence our model. However, we may have missed other important variables or mischaracterised the causal link between the variable and adoption. We used several approaches to support our assumption. First, the conceptual model draws on Diffusion of Innovation theory, a well-established theoretical framework used across diverse contexts, including in conservation research [e.g., 13-15, 17, 18, 19, 54]. In general, using a robust theoretical framework in empirical modelling can reduce the chance of spurious results by grounding the analysis in established concepts and principles [55, 56]. Second, our analysis was a transdisciplinary collaboration between university scientists and FLMMA network practitioners in Fiji. This collaboration provided a unique, in-depth view of the processes influencing adoption and how they can be mathematically represented.

While our modelling framework has been built around a specific case study, our agent-based model can readily be adapted to other data sets. To fit the spatiotemporal adoption pattern in another data set, one merely needs to provide the associated properties (***X*** in eq. 1) of the agents (here, villages) and specify the networks between them (e.g., *N* in eq. 1). We thus plan to apply our modelling framework to a broader set of case studies in the future and encourage others to adapt our code to theirs.

In conclusion, predictive conservation science approaches may help guide efforts to tackle the triple challenge of reversing biodiversity loss, halting climate change, and supporting human well-being. We provide one of the first studies to operationalize Diffusion of Innovations theory in an agent-based model to forecast the adoption and spread of conservation actions. We find that accurately forecasting how, where, and when community conservation practices will likely be adopted requires detailed and specific knowledge about the communities involved. However, well-designed models that build on this context-specific understanding can help forecast how changes in the characteristics of a system alter the patterns of adoption and spread of conservation measures. Such models can be integrated into adaptive management systems to track progress and course-correct to scaling targets. The models can also help anticipate where and among which social groups the socio-ecological benefits and costs of conservation actions will occur. Furthermore, insights from these models could be combined with effectiveness evidence to predict which measures are likely to deliver impact at scale 33]. To support this goal, we provide a step-by-step framework that other researchers could use to understand, forecast, and inform the scaling of community-oriented conservation measures across contexts.

## Acknowledgements

We thank the Turner-Kirk Trust for supporting this research. MM, TP, and MC thank the Leverhulme Trust for the research grant (RPG-2021-440). ACSJ is supported by the Eric and Wendy Schmidt AI in Science Postdoctoral Fellowship, a Schmidt Sciences program.

## Data and code availability statement

Summary data supporting this study’s findings are included in the paper and its Supplementary Information. FLMMA member village data are available upon request from the FLMMA Secretariat (email contact). Covariate data used for matching were provided by the Ministry of Lands and Mineral Resources and Fiji Roads Authority, with the exception of coral cover data, which are publicly available from the Millennium Coral Reef Mapping Project (available at https://oceancolor.gsfc.nasa.gov/cgi/landsat.pl). Raw data from the interviews are available on request from the corresponding author (MM) with reasonable restrictions, as respondents belong to the Indigenous iTaukei group and have additional protections under our ethical review process. Data used for the analysis will be made available no more than 2 weeks after the data use agreement has been agreed and signed. Code and illustrative synthetic data are available at: https://github.com/ASoelvsten/COALA.

## A Supplementary Table

**Table 1:** Table of parameters and the details of the associated survey data. All information is available for the 122 villages in the data set discussed in this paper.

| Explanation | Parameter | Data details |
| --- | --- | --- |
| Presence of support organisations | $\beta_1$ | Whether organisations or government departments currently support the qoliqoli (binary) |
| Compatibility with needs | $\beta_2$ | The degree to which LMMAs are seen as consistent with the existing needs of potential adopters (5 point likert item) |
| Supportive national policies | $\beta_3$ | The extent to which government oppose or support the community in developing their own rules about the marine resources within [usual fishing grounds]? (5 point likert item) |
| Observable benefits | $\beta_4$ | The extent to which the result of having a resource management is visible to everyone (5 point likert item) |
| Overall benefits | $\beta_5$ | Whether the community has benefited from the qoliqoli rules or management of marine resources in the qoliqoli over the past ten years? (5 point likert item) |
| Political empowerment (presence of champions) | $\beta_6$ | Whether the community champions who support resource management within the community (5 point likert item) |
| Access to decision-making structures (district level) | $\beta_7$ | Positive responses to “is there a Yaubula Committee at the district level?” (binary) |
| Supportive institutions | $\beta_{10}$ | Whether the people in this village generally trust the government (5 point likert item) |
| Perceived knowledge (coral reefs) | $\beta_{13}$ | Whether marine resource users in [group represented] are very knowledgeable about what affects the health of the coral reefs (5 point likert item) |
| Perceived knowledge (quantity of fish) | $\beta_{14}$ | Whether marine resource users in [group represented] are very knowledgeable what affects the quantity of fish and shellfish in the qoliqoli (5 point likert item) |
| Perceived knowledge (diversity of fish) | $\beta_{15}$ | Whether marine resource users in [group represented] are very knowledgeable the diversity of fish and shellfish in the qoliqoli (5 point likert item) |
| Fairness of decision-making processes | $\beta_{16}$ | Thinking about the way in which decisions are made in the village about using and managing the qoliqoli: how fair is the decision-making process in your opinion? (5 point likert item) |
| Fairness of rules | $\beta_{17}$ | Thinking about the way in which decisions are made in the village about using and managing the qoliqoli: How fair would you say that the actual rules about using the qoliqoli are?” (5 point likert item) |

## B Supplementary Figures

**Figure S1:**
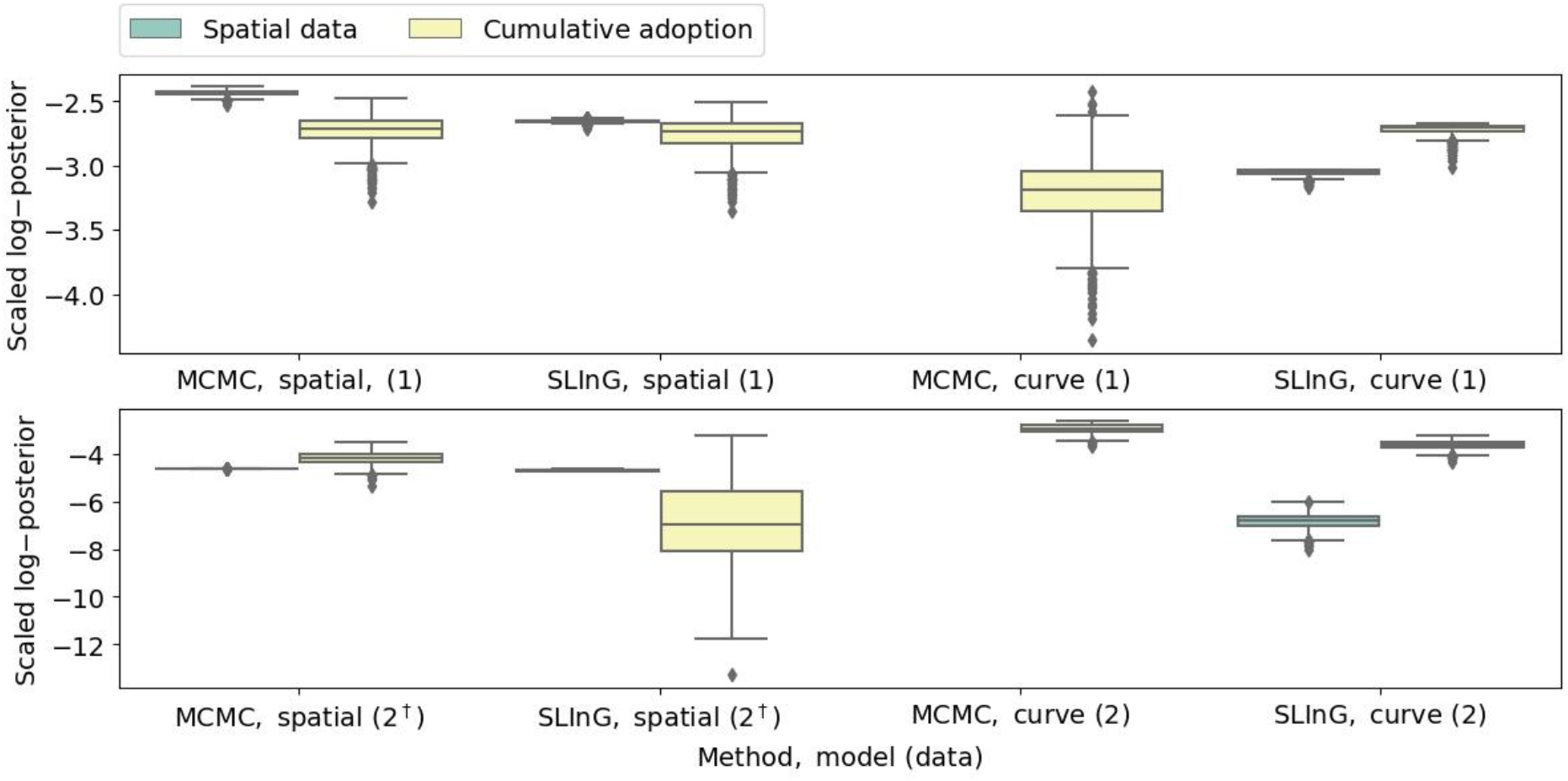
Performance metric for a subset of the fits presented in the results section. The plot presents results obtained using the MCMC ensemble sampler as well as the sparse sampling algorithm SLInG. Simulations were computed fitting only the curve summarising the total number of adopters as a function of time or the entire spatial distribution of adopters. This is specified on the horizontal axes. The upper panel relates to the data set (1) with 122 villages, while the lower panel relates to the larger data set (2) with 627 villages with (dagger) and without the inclusion of β_11_ and β_12_. The metric is proportional to the total log-probability assigned to the parameter set when matching the adoption by the listed data set (either by individual villages in space and time or performing a comparison to the total number of adopters as a function of time. This is specified in the legend). When the log-probability obtained from including the spatial is not shown, the corresponding probability is zero for all or most samples. The metrics have been scaled to make them comparable within each panel.

**Figure S2:**
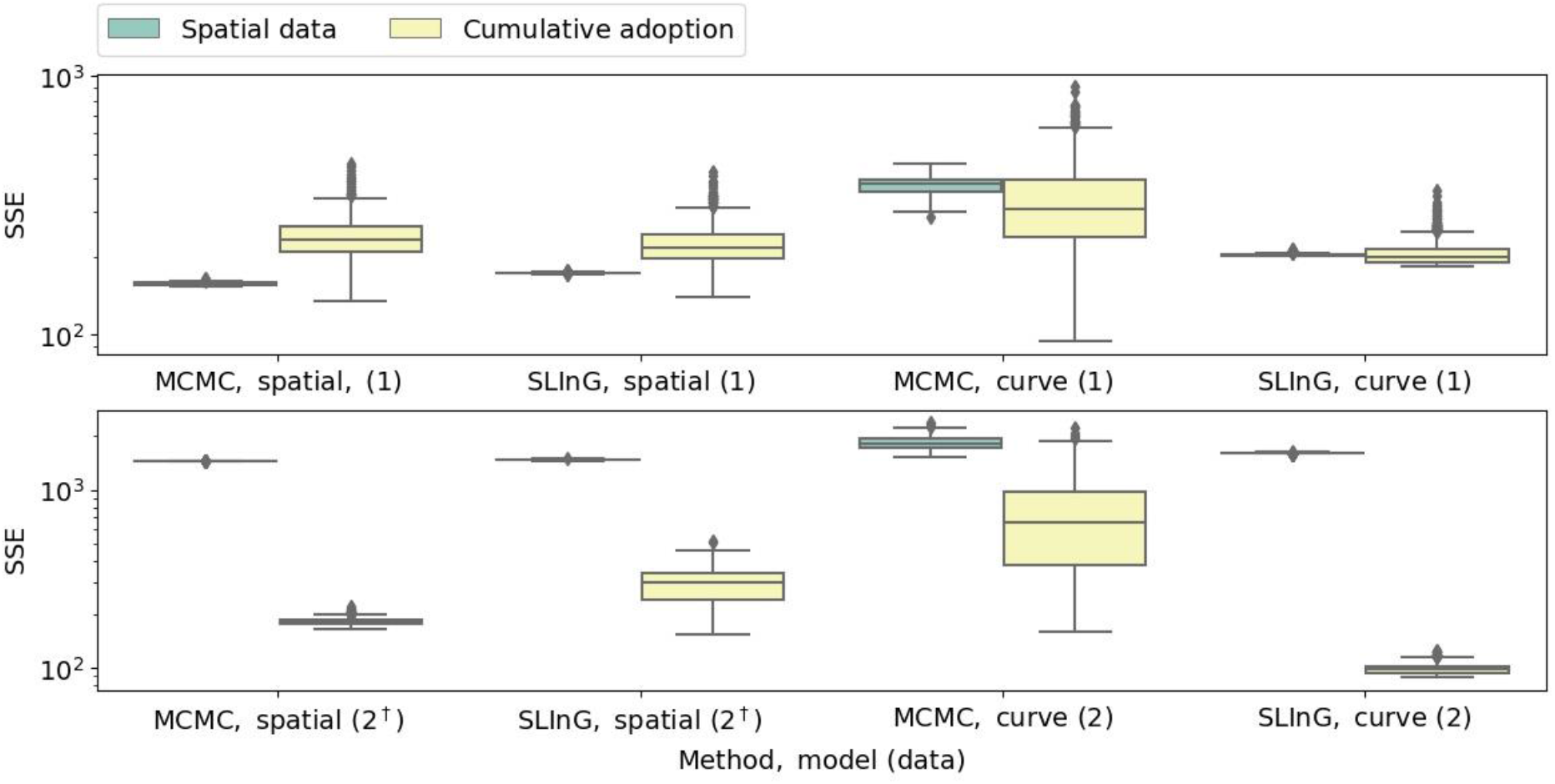
Pendant to Figure S1 showing the sum of squared differences between the predicted probabilities (on a scale from zero to one) and the observed adoption status: non-adopters (0) vs. adopters (1).

**Figure S3:**
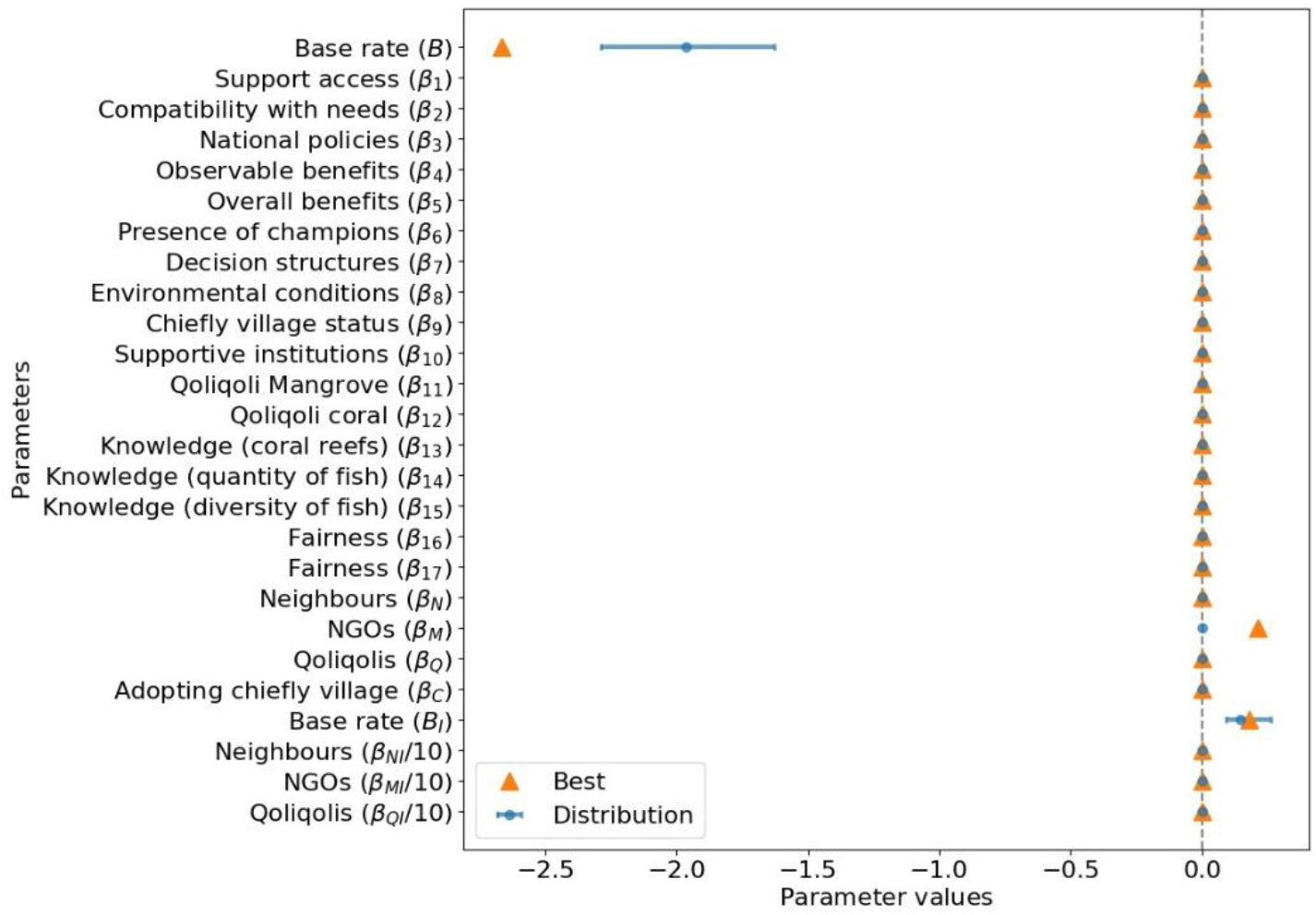
Pendant to Figure 3 obtained using SLInG fitting the total number of adopters as a function of time for the data set with 122 villages for which information on all perceived benefits is available. Note that the best-fitting parameter set, i.e., the parameter set that leads to the lowest error, within the Markov chain (including parts of the burning) might deviate from the obtained posterior distribution since the sparsity inducing priors favour solutions with fewer parameters and lower parameter values, although this is not the case here.

**Figure S4:**
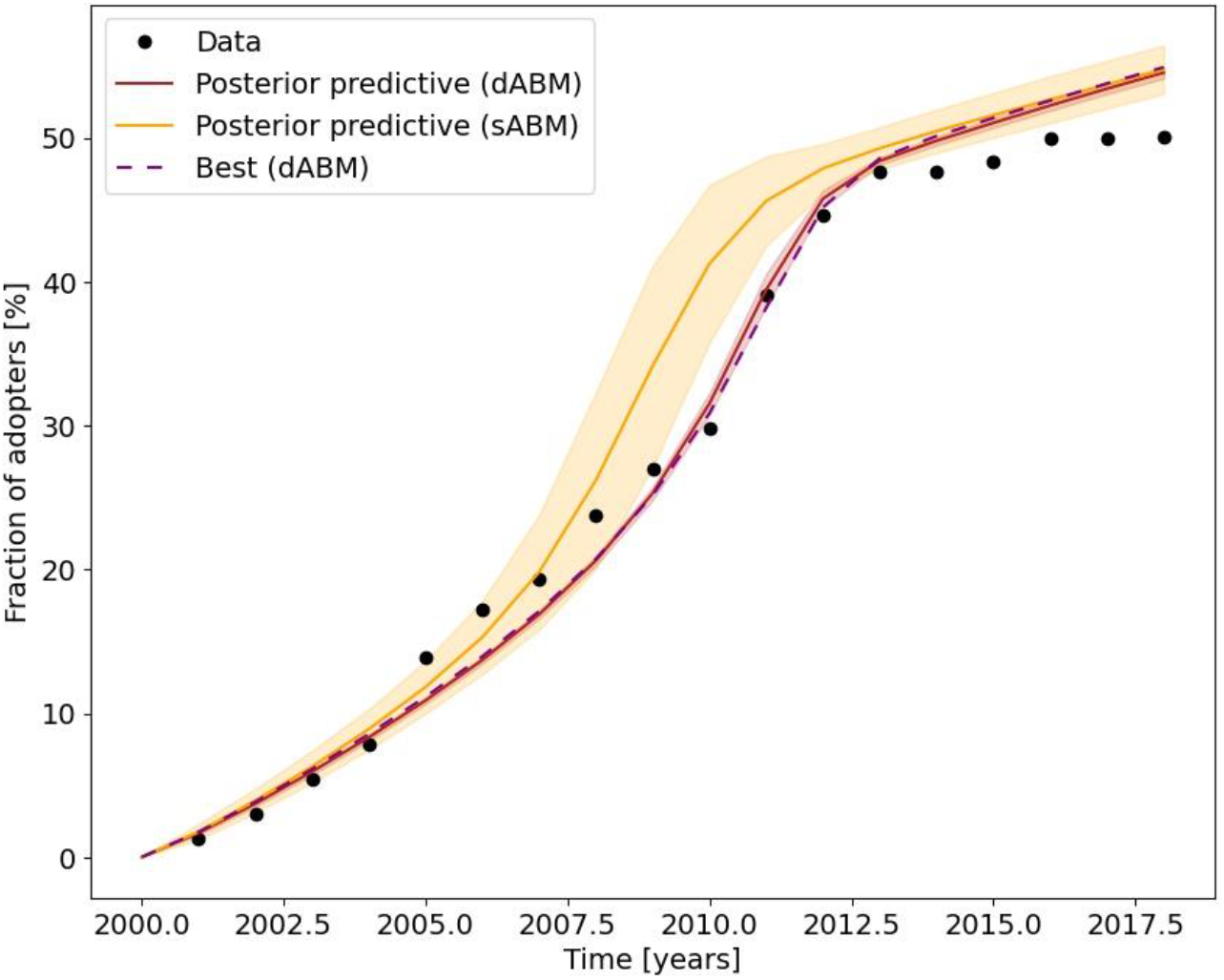
Adoption as a function of time for the large data set containing 627 villages. The fit includes only B, β_N_, β_M_, β_Q_, B_I_, β_NI_, β_MI_, and β_QI_. It was obtained with SLInG using sparsity-inducing priors for all parameters except B and B_I_. The shaded area signifies 95 per cent confidence intervals while the solid line shows the corresponding median. The dashed line shows predictions by the best parameter set among models in the Markov chains.

**Figure S5:**
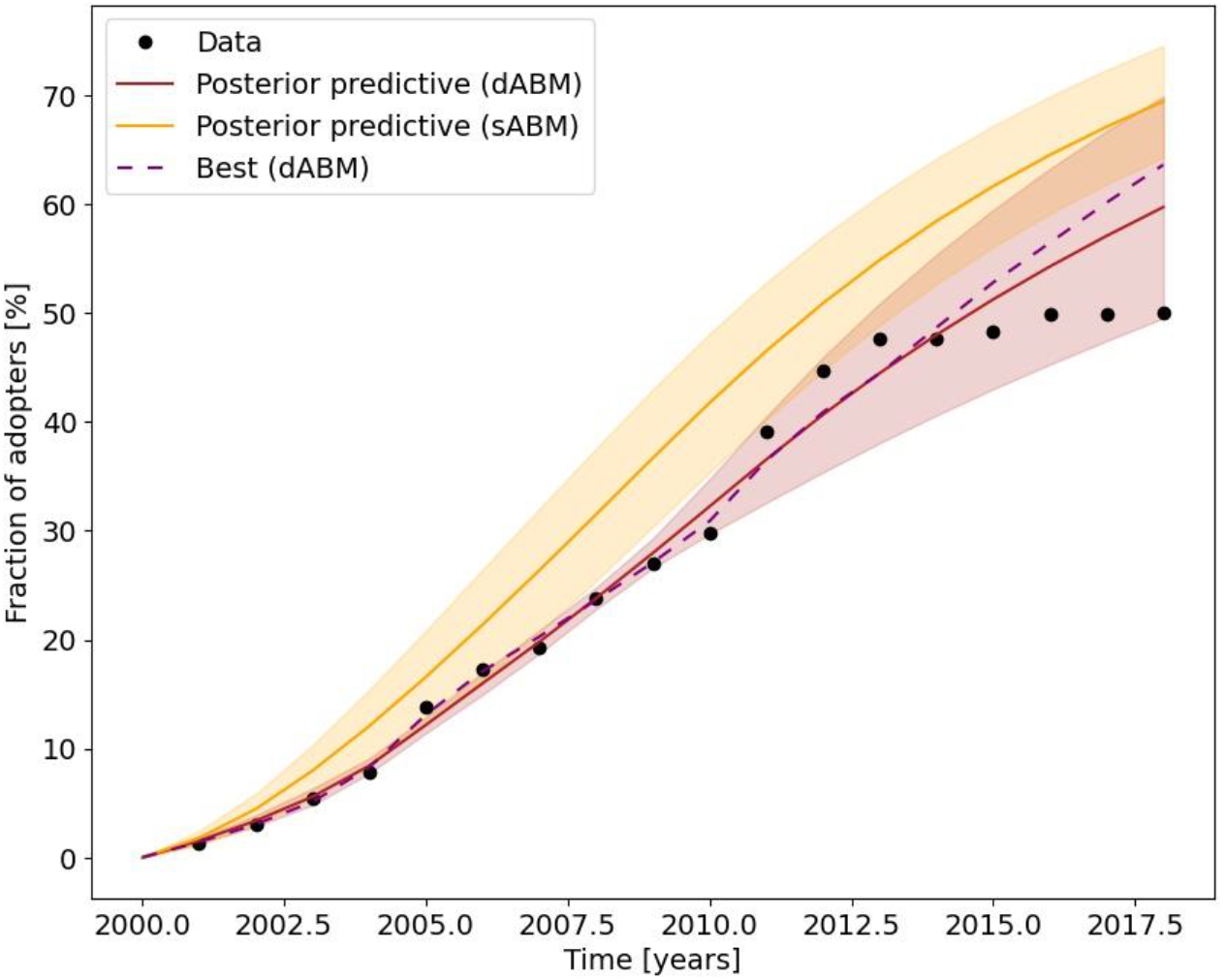
Pendant to Figure S4 for the data set with 627 villages using the MCMC ensemble sampler. The error bars indicate 95 per cent confidence intervals. The solid line corresponds to the median of the posterior distribution. The fit does not include β_11_ and β_12_.

**Figure S6:**
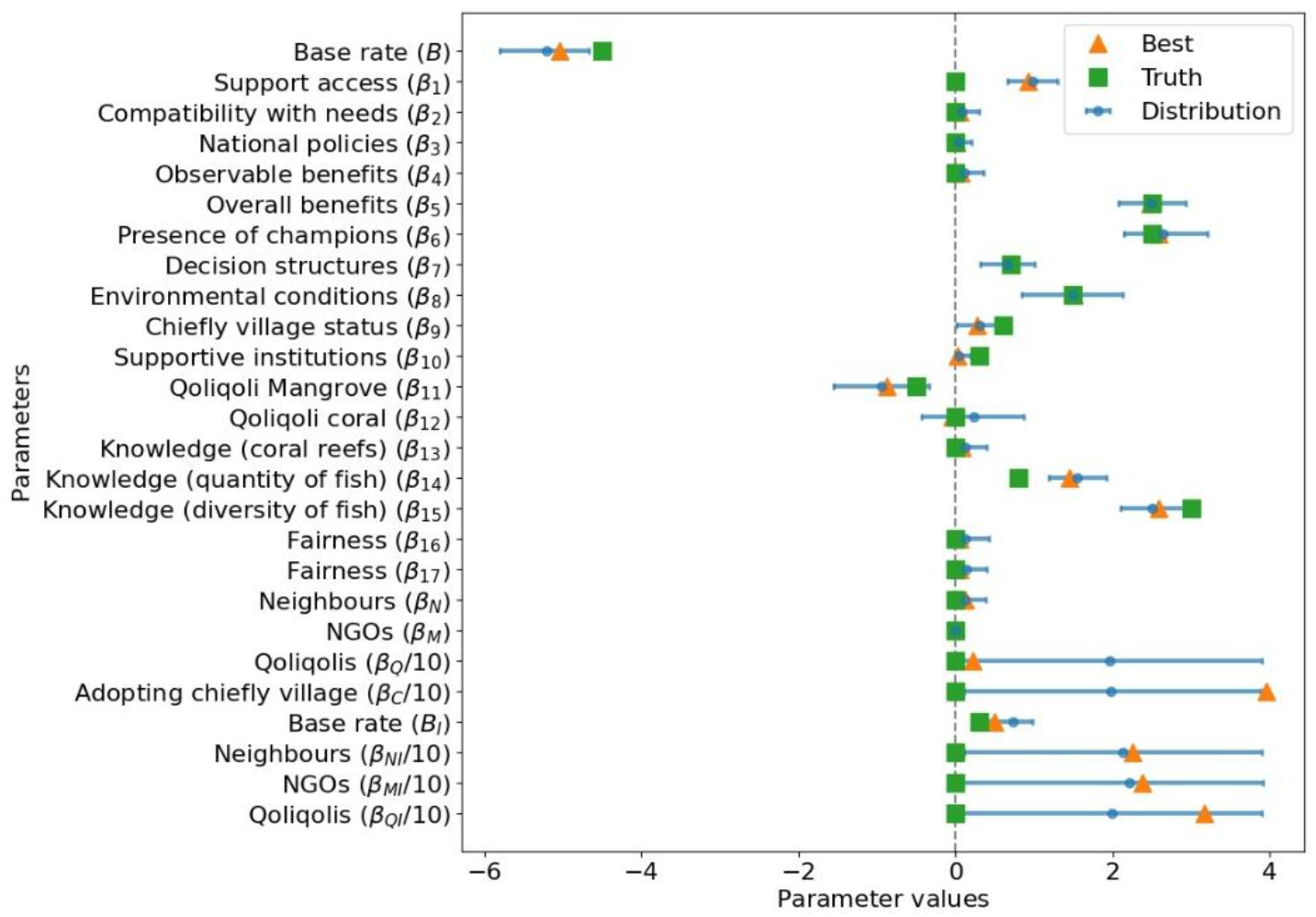
Results from self-consistent test. We created a synthetic dataset including 128 villages using the stochastic agent-based model. We then inferred the underlying parameter setting using the MCMC ensemble sampler and the deterministic model. Comparing the inferred parameter values with the true parameter values of the synthetic data set, we see that our framework correctly determined the sign and magnitude of most parameters despite the low number of villages and the spurious effects that a small dataset of this type might present.

## C: Second data set

In addition to the data set presented in the main text, we analysed a larger data set containing 627 villages. For these villages, information was available on all three networks through which the agents in our model communicate (*N_i_*, *M_i_* and *Q_i_*) and for the perceived benefits associated with β11 and β12. Just like for the smaller data set, we find that we recover the overall trend of the curve describing the total number of adopters with time when we fit the spatiotemporal adoption pattern using the MCMC ensemble sampler or SLInG. Indeed, when imposing sparsity-inducing priors, we consistently find that the base rates take non-zero values whether or not we include β_11_ and β_12_ as free parameters.

When we fit the total number of adopters as a function of time for the data set containing 627 villages using the MCMC ensemble sampler, we obtain a decent fit to data if we do not include β_11_ and β_12_ (cf. Figure S5). However, if we include these parameters, we can no longer constrain β_N_ and β_Q_. Indeed, we merely recover the priors for β_11_, β_N_ and β_Q_, which significantly increases the uncertainties on the predicted number of adopters.

When we repeat the analysis using SLInG, only B, B_I_ and β_M_ take non-zero values. The results of this fit are shown in Figure S4. Again, we note that this is not to say that the supporting organisations play a more vital role than either local neighbours or other connections but rather that a single social network is sufficient to recover the cumulative pattern of adoption.

